# ENIGMA: An Enterotype-Like Unigram Mixture Model for Microbial Association Analysis

**DOI:** 10.1101/397091

**Authors:** Ko Abe, Masaaki Hirayama, Kinji Ohno, Teppei Shimamura

## Abstract

**Background:** One of the major challenges in microbial studies is to discover associations between microbial communities and a specific disease. A specialized feature of microbiome count data is that intestinal bacterial communities have clusters reffered as enterotype characterized by differences in specific bacterial taxa, which makes it difficult to analyze these data under health and disease conditions. Traditional probabilistic modeling cannot distinguish dysbiosis of interest with the individual differences.

**Results:** We propose a new probabilistic model, called ENIGMA (Enterotype-like uNIGram mixture model for Microbial Association analysis), to address these problems. ENIGMA enables us to simultaneously estimate enterotype-like clusters characterized by the abundances of signature bacterial genera and environmental effects associated with the disease.

**Conclusion:** We illustrate the performance of the proposed method both through the simulation and clinical data analysis. ENIGMA is implemented with R and is available from GitHub (https://github.com/abikoushi/enigma).

## Introduction

More than 100 trillion microbes live on and within human beings and consists of complex microbial communities (microbiota). The majority of microbes cannot be cultured in laboratories, which makes it difficult to understand which individual microorganisms mediate vital microbiome-host interactions under health and disease conditions. However, recent important advances in high-throughput sequencing technology have allowed us to observe the composition of these intestinal microbes. That is, for each sample drawn from an ecosystem, the number of occurrences of each operational taxonomic units (OTUs) is measured and the resulting OTU abundance are summarized at any level of the bacterial phylogeny. Discovering recurrent microbial compositional patterns that are related with a specific disease is a significant challenge since individuals with the same disease typically harbor different microbial community structures.

The recent large-scale sequencing surveys of the human intestinal microbiome, such as the US NIH Human Microbiome Project (HMP) and the European Metagenomics of the Human Intestinal Tract project (MetaHIT), have shown considerable variations in microbiota composition among individuals [1, 2]. In particular, the presence of community clusters characterized by differences in the abundance of signature taxa, referred to as enterotypes, have been first reported in humans [3]. Later, other studies found enterotype-like clusters which might reflect features of host-microbial physiology and homeostasis in different species [4, 5] or across human body sites [6–9]. These observed microbial stratification has motivated the development of methods to examine unknown clusters of microbial communities.

Probabilistic modeling of microbial metagenomics data often provides a powerful framework to characterize the microbial community structures [10–12]. For example, Knights *et al.* [10] applied a Dirichlet prior to a single-level hierarchy and proposed a Bayesian approach to estimate the proportion of microbial communities. Holmes *et al.* [11] extended the Dirichlet prior to Dirichlet multinomial mixtures to facilitate clustering of microbiome samples. Shafiei *et al.* [12] proposed a hierarchical model for Bayesian inference of microbial communities (BioMiCo) to identify clusters of OTUs related with environmental factors of interest.

However, such models are not suitable for identification of enterotype-like clusters of microbial communities doe to the following two reasons. First, the frameworks of Knights *et al.* [10] and Holmes *et al.* [11] do not explicitly address the association between the microbial compositional patterns and environmental factors of interest. Second, the framework of Shafiei *et al.* [12] models the structure of each sample by a hierarchical mixture of multinomial distributions that are dependent to factors of interest. It is known that individual host properties such as body mass index, age, or gender cannot explain the observed enterotypes [3]. Thus, such enterotype-like clusters that describes interindividual variability among humans do not always to directly affect host probabilities such as diseases ranging from localized gastroenterologic disorders to neurologic, respiratory, metabolic hepatic, and cardiovascular illnesses.

Here, we introduce a novel probabilistic model of a microbial community structures, called ENIGMA (Enterotype-like uNIGram mixture model for Microbial Association analysis), to address these problems. ENIGMA includes the following contributions:

1. ENIGMA takes OTU abundances as input and models each sample by underlying unigram mixture whose parameters are represented by unknown group effects and known effects of interest. The group effects are represented by the baseline parameters which change with a latent group of microbial communities. One of the most important features for our model is that the group effects are independent of the effects of interest. This enables to separate interindividual variability and fixed effects of the host properties related with disease risk.
2. ENIGMA is regarded as a Bayesian learning for the association between community structure and factors of interest. Our model can be used to simultaneously learn how enterotype-like clusters of OTUs contributes to microbial structure and how microbial compositional patterns might be related to the known features of the sample.
3. We provide an efficient learning procedure for ENIGMA by using a Laplace approximation to integrate out the latent variables and estimate the evidence of the complete model and the credible intervals of the parameters. The software package that implements ENIGMA in the R environment is available from https://github.com/abikoushi/enigma.

We describe our proposed framework and algorithm in section named “Methods”. We evaluate the performance of ENIGMA on simulated data in terms of its accuracy to estimate parameters and identify clusters in Section named “Simulation Study”. We apply ENIGMA to clinical metagenomics data and demonstrate how ENIGMA simultaneously identifies enterotype-like clusters and gut microbiota related with Parkinson’s disease (PD) in Section named “Results on Clinical Data”.

## Methods

Suppose that we observe microbiome count data of *K* taxa for *N* samples with *M* individual host properties, (*y*_*nk*_, *x*_*nm*_) (*n* = 1,…, *n*; *k* = 1,…, *K*; *m* = 1,…, *M*) where 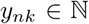 represents the abundance of the *k*-th taxa in the *n*-th sample and *x*_*nm*_ represents a binary variable such that *x*_*nm*_ = 1 if the *n*-th sample has the *m*-th host property and *x*_*nm*_ = 0 otherwise. Here the word “taxa” could be at any level of the bactgerial phylogeny, e.g., species, genes, family, order, etc.

### Model

Figure 1 illustrates the plate diagram of the proposed model for metagenome sequencing, where ***y****_n_* is the read count vector of the *n*-th sample, ***x****_n_* is the vector of the host properties of the *n*-th sample and *z*_*n*_ ∈ {1,…, *L*} is a latent class of the *n*-th sample. Our model is a simple extension of unigram mixture model. We assume that each sample is generated from a multinomial distribution with the parameter vector ***p****_n_* = (*p*_*n*1_,…, *p*_*nK*_)^*T*^. The elements of ***p***_*n*_, *p*_*nk*_(*k* = 1,…, *K*) are probabilities of the occurrence of the *K* taxa for the *n*-th sample. We also assume that *p*_*nk*_ can be influenced independently by the environmental factor on the taxa that is common to all latent classes and the interindividual factor on the latent enterotype-like classes. More specifically, the generative process of ENIGMA is defined by:
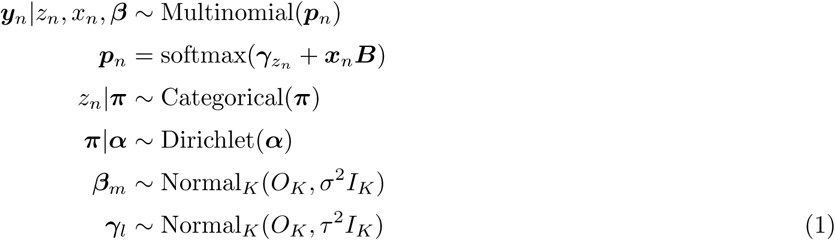
where ***γ****_l_* is baseline parameter (*K*-dimensional vector) which change with the latent class, *M × K* matrix ***B*** = (*β*_*mk*_) is effect of a environmental factor common the all enterotypes, ***β****_m_* is a *m*-th row-vector of ***B, π*** = (*π*_1_,…, *π*_*L*_) is a mixing ratio of components, *O*_*K*_ is *K*-dimensional zero matrix and *I*_*K*_ is *K*-dimensional identity matrix. Here softmax function is defined by softmax 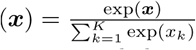 for a vector ***x*** = (*x*_1_,…, *x*_*K*_)^*T*^ using element-wise exponential function and the probability function of categorical distribution is parameterized as Pr(*z* = *l|****π***) = *π*_l_, *l* ∈ {1,…, *L*}. In a Bayesian approach we need to define prior distributions for ***π, β***, and ***γ***_*l*_. We set a prior based on the Dirichlet distribution for ***π***, and flat priors to the hyperparameters *σ* and *τ* for ***β*** and ***γ***, respectively. For the convenience of later section, let ***p****′_l_* = softmax(***γ***_*l*_) be probabilities of the occurrence of bacteria in the latent classes *l*.

**Figure 1.**
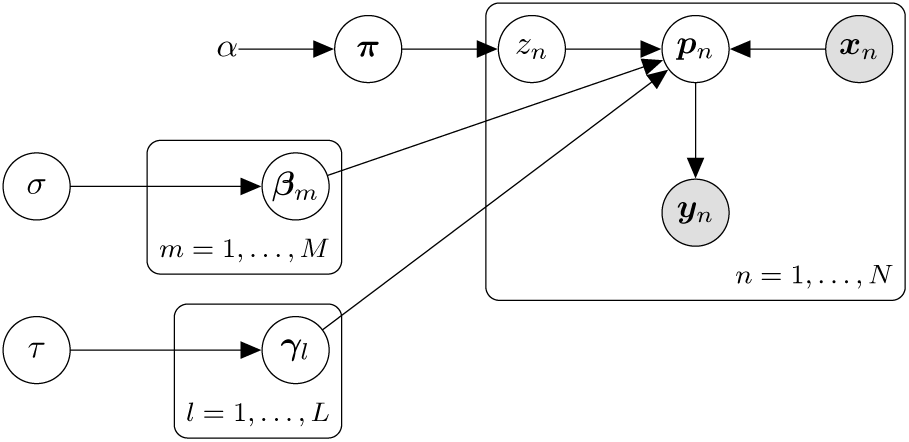
Plate diagram of the model for ENIGMA. *yn* is affected from environmental factors ***x****n* and latent variables ***z****n*.

### Parameter estimation

Let us denote observed matrix by ***Y*** = (*y*_*nk*_), ***X*** = (*x*_*nm*_), the unknown parameters by ***θ*** = (***α, B, γ***_1_,…, ***γ****_L_, σ, τ*) and their prior by *φ*(***θ***). The posterior distribution is represented as follows:
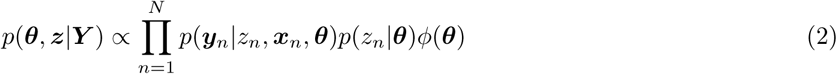
First, latent variable *z*_*n*_ must be marginalized. The likelihood belongs to
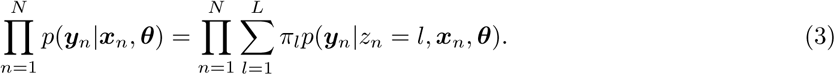
The posterior distribution is proportional to product of the likelihood and prior density:
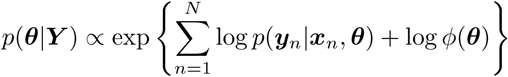
Let 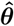 be the MAP estimator of ***θ***, found by maximizing log *p*(***θ, Y, X***).

We use a Laplace approximation [15] for parameter estimation. A Taylor expansion around 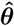 gives
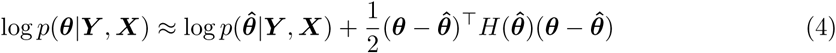
where and *H*(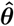) is Hessian of log *p*(***θ***|***Y, X***) evaluated at 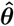. Eq.4 gives
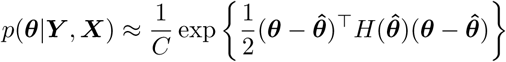
where *C* is normalizing constant. This relation shows that *p*(*θ|****Y, X***) can be approximated by normal distribution *N* (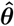, *H*^−1^(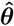)). Credible intervals can be calculated from this multivariate normal distribution.

We used stochastic programming language Stan (http://mc-stan.org/) for its implementation. The MAP estimators were obtained by L-BFGS method. Credible intervals were computed from the using a Stan function to compute the Hessian at the MAP estimates.

After fitting the model, we are left with the task of classify the enterotype of each samples. The conditional probability of *z*_*n*_ = *l* is
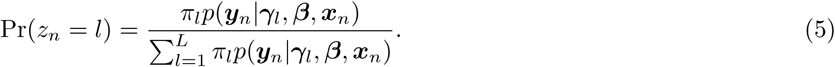
This is the probability which *n*-th sample belong enterotype *l*. Then, *n*-th sample is then classified into the *l*-th enterotype that maximizizes the conditional probability gven by Eq.5.

### Model Selection

We are also interested in whether or not the whole set rather than individual bacteria is related to the environmental factors of interest. We consider the comparison between the two models when ***B*** /= **0** and ***B*** = **0**. We can use the log marginal likelihood as the goodness of fit for model comparison. The marginal likelihood is given by
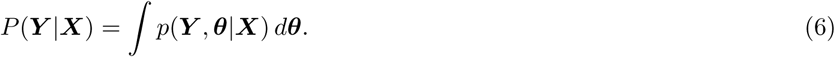
From Eq.4, we have
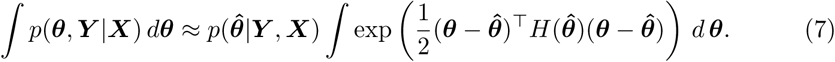
So, log marginal likelihood is approximated by following formula:
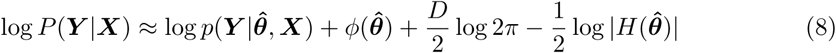
where *D* is the number of free parameters. In model comparison, we choose the model with the larger log marginal likelihood.

## Simulation Study

To show the performance of ENIGMA, we conducted several experiments by simulation. The synthetic data can be naturally produced via our generative process given by Eq.1. We set *M* = 1, *L* = 3, *π*_*l*_ = 1/3, and ***α*** = (1, 1, 1)^*T*^. We first generated ***B*** and ***γ****_l_* from the standard normal distribution. The variables ***x****_n_, z_n_*, and ***y****_n_* are then sampled from the Bernoulli distribution with probability of 0.5, the categorical distribution, and the multinomial distribution, respectively. For the above parameter setting, we randomly generate a count dataset of 100 taxa for 100 samples for evaluation.

- **Coverage probability (CP)**: The coverage probability is the proportion of the time that the interval contains the true value. A discrepancy between the coverage probability and the nominal coverage probability frequently occurs. When the actual coverage is greater than the nominal coverage, the interval is called conservative. If the interval is conservative, there is no inconsistency in interpretation.
- **Bias**: The bias of ***B*** is defined by difference between true value and estimated value *E*[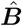] *−* ***B***.
- **Standard error (SE)**: The standard error is the standard deviation of the estimate. The smaller standard error indicates the higher accuracy of estimation.
- **Root mean squared error (RMSE)**: The RMSE is defined by 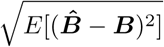. The smaller RMSE indicates the higher accuracy of estimation.
- **Accuracy**: The accuracy is the percentage of samples correctly classified into original group.

For calculating these metrics, we note that we calculated the sample means and standard deviations of 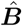 and (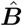 *−* ***B***)^2^ from the 10,000 synthetic datasets.

Figure 2 shows the comparison of true ***B*** and the mean and standard deviation of estimates 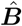 through the 10,000 simulations. We observed that the points are arranged diagonally, which implies the estimator of ENIGMA is unbiased. We also calculated the proportion of the time that the 95% credible interval contains the true value of ***B***. We found that this proportion is greater than nominal value 0.95 for all ***B*** in Figure 3. Table 2 shows the coverage probability (CP), bias, standard error (SE), and RMSE of 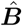, respectively. We observed that the bias and standard error decrease when *β*_*mk*_ is large (i.e. the corresponding abundance is large). We also found that the accuracy of classification given by Eq.5 is exactly 100%. Thus, these results indicate that ENIGMA can produce reasonable estimates.

**Figure 2.**
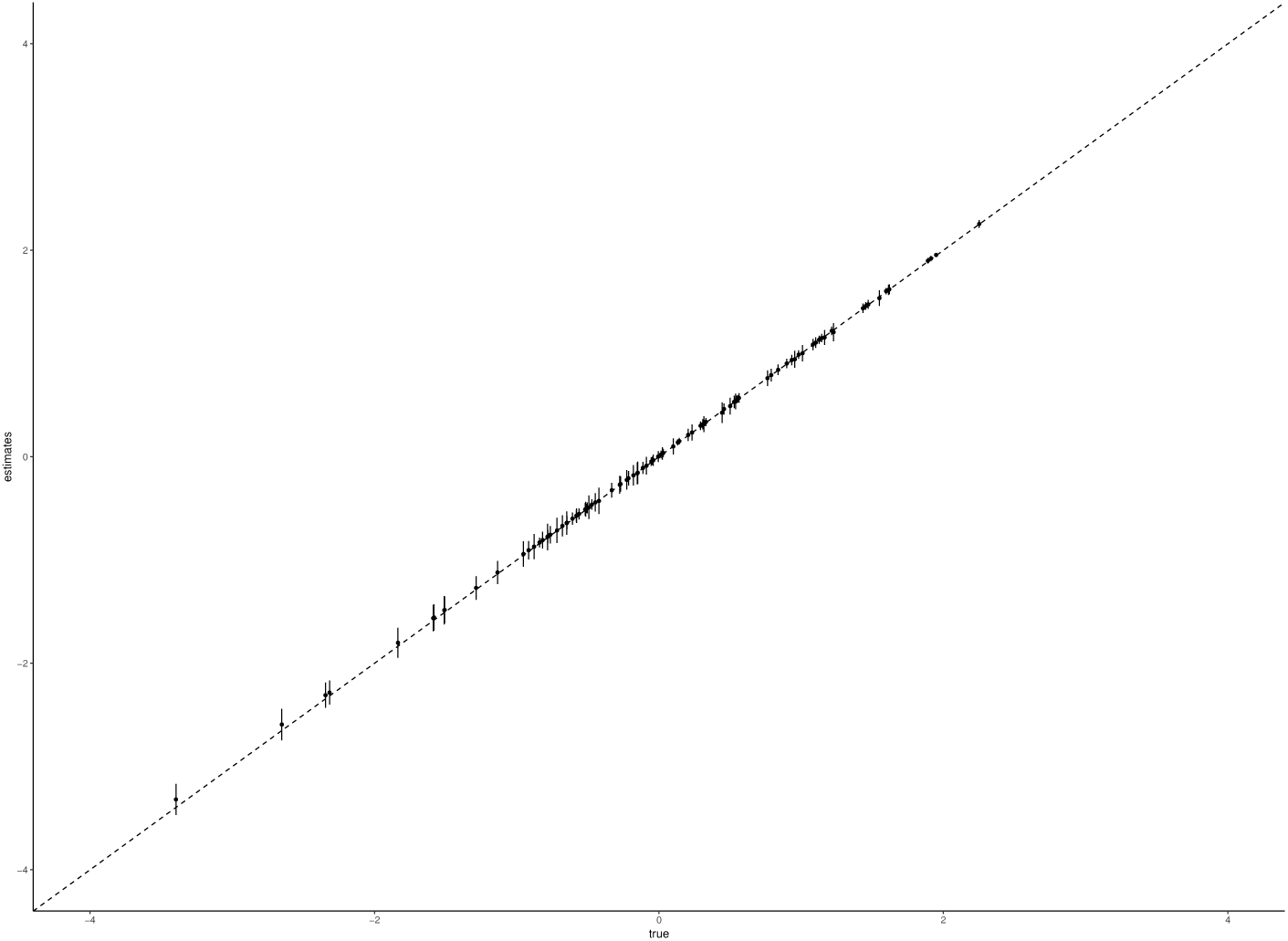
Simulation result of *B*. The comparison true ***B*** and the mean of ***B*ˆ**. The error bars indicates SE.

**Figure 3.**
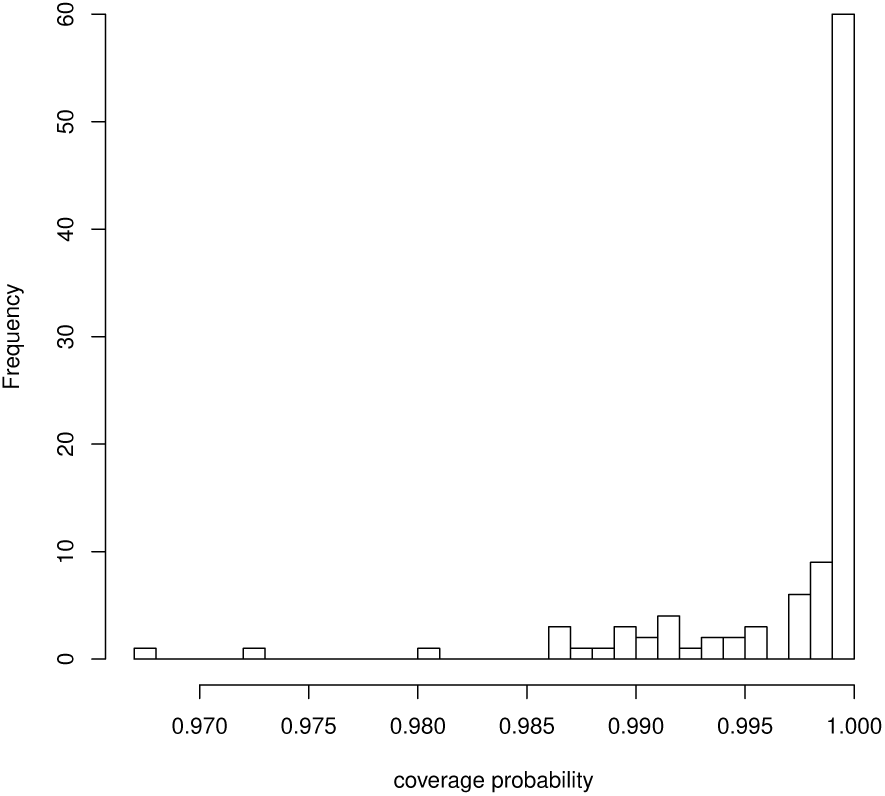
Coverage probability of *B*. The histogram of coverage probability of ***B***.

## Results on Clinical Data

To validate the performance of ENIGMA on discovering clusters of micribial communities and associations between microbes and a specific disease, we applied ENIGMA to the real metagenomic sequencing data from Scheperjans *et al.* [16], Hill-Burns *et al.* [17], Heintz-Buschart *et al.* [18] and Hopfner *et al.* [19]. The data is analized by sequencing the bacterial 16S ribosomal RNA genes sampled from patients of Parkinson’s disease (PD) and control in Finland, USA, and Germany. Table 1 shows the summary statistics of the data. The OTUs are mapped to the SILVA taxonomic reference, version 132 (https://www.arb-silva.de/) and the abundances of family-level taxa are calculated. Following the evidence of Arumugam *et al.* [3], the number of latent classes in ENIGMA is chosen to be *L* = 3. We set the hyperparameters of Dirichlet prior ***α*** = (1, 1, 1)^*T*^, which is equivalent to a non-informative prior.

**Table 1.** The data summary

|  | PD | CO |
| --- | --- | --- |
| Finland | 74 | 74 |
| German | 55 | 64 |
| USA | 207 | 139 |

**Table 2.** Coverage probability (CP), bias, standard error (SW) and RMSE of ***B*ˆ**

| $\beta$ | CP | bias | SE | RMSE | $\beta$ | CP | bias | SE | RMSE |
| --- | --- | --- | --- | --- | --- | --- | --- | --- | --- |
| -3.40 | 0.97 | 0.08 | 0.15 | 0.17 | -0.04 | 1.00 | 0.01 | 0.05 | 0.05 |
| -2.65 | 0.97 | 0.06 | 0.15 | 0.16 | -0.04 | 1.00 | 0.01 | 0.05 | 0.05 |
| -2.34 | 0.99 | 0.04 | 0.12 | 0.13 | -0.01 | 1.00 | 0.01 | 0.05 | 0.05 |
| -2.32 | 0.99 | 0.03 | 0.12 | 0.12 | 0.01 | 1.00 | 0.01 | 0.04 | 0.04 |
| -1.83 | 0.98 | 0.03 | 0.14 | 0.15 | 0.02 | 1.00 | 0.01 | 0.06 | 0.06 |
| -1.59 | 0.99 | 0.02 | 0.13 | 0.13 | 0.02 | 1.00 | 0.01 | 0.04 | 0.05 |
| -1.58 | 0.99 | 0.03 | 0.13 | 0.13 | 0.03 | 1.00 | 0.01 | 0.04 | 0.04 |
| -1.51 | 0.99 | 0.02 | 0.14 | 0.14 | 0.10 | 1.00 | -0.00 | 0.08 | 0.08 |
| -1.51 | 0.99 | 0.02 | 0.13 | 0.13 | 0.13 | 1.00 | 0.01 | 0.03 | 0.03 |
| -1.29 | 0.99 | 0.02 | 0.11 | 0.11 | 0.14 | 1.00 | 0.01 | 0.03 | 0.03 |
| -1.14 | 0.99 | 0.01 | 0.11 | 0.11 | 0.21 | 1.00 | 0.01 | 0.06 | 0.06 |
| -0.95 | 1.00 | 0.01 | 0.09 | 0.09 | 0.23 | 1.00 | 0.00 | 0.08 | 0.08 |
| -0.95 | 0.99 | 0.01 | 0.12 | 0.12 | 0.29 | 1.00 | 0.01 | 0.04 | 0.04 |
| -0.92 | 1.00 | 0.01 | 0.09 | 0.09 | 0.31 | 1.00 | 0.01 | 0.05 | 0.05 |
| -0.88 | 0.99 | 0.01 | 0.12 | 0.12 | 0.32 | 1.00 | 0.00 | 0.08 | 0.08 |
| -0.84 | 1.00 | 0.01 | 0.05 | 0.05 | 0.33 | 1.00 | 0.01 | 0.04 | 0.04 |
| -0.82 | 1.00 | 0.01 | 0.08 | 0.08 | 0.44 | 0.99 | -0.02 | 0.10 | 0.10 |
| -0.78 | 0.99 | 0.01 | 0.13 | 0.13 | 0.46 | 1.00 | 0.01 | 0.05 | 0.05 |
| -0.78 | 1.00 | 0.01 | 0.07 | 0.07 | 0.50 | 1.00 | -0.01 | 0.08 | 0.08 |
| -0.76 | 1.00 | 0.01 | 0.08 | 0.08 | 0.53 | 1.00 | 0.00 | 0.06 | 0.06 |
| -0.72 | 0.99 | 0.00 | 0.12 | 0.12 | 0.54 | 1.00 | -0.00 | 0.08 | 0.08 |
| -0.68 | 1.00 | 0.01 | 0.10 | 0.10 | 0.55 | 1.00 | 0.01 | 0.04 | 0.04 |
| -0.65 | 0.99 | 0.01 | 0.11 | 0.11 | 0.55 | 1.00 | 0.01 | 0.03 | 0.03 |
| -0.65 | 0.99 | 0.01 | 0.11 | 0.11 | 0.56 | 1.00 | 0.01 | 0.05 | 0.05 |
| -0.65 | 1.00 | 0.01 | 0.06 | 0.06 | 0.76 | 1.00 | -0.00 | 0.07 | 0.07 |
| -0.61 | 1.00 | 0.01 | 0.06 | 0.06 | 0.79 | 1.00 | 0.00 | 0.06 | 0.06 |
| -0.58 | 1.00 | 0.01 | 0.06 | 0.06 | 0.84 | 1.00 | 0.00 | 0.05 | 0.05 |
| -0.58 | 1.00 | 0.01 | 0.07 | 0.07 | 0.90 | 1.00 | 0.01 | 0.04 | 0.04 |
| -0.56 | 1.00 | 0.01 | 0.05 | 0.05 | 0.93 | 1.00 | 0.00 | 0.05 | 0.05 |
| -0.52 | 1.00 | 0.01 | 0.06 | 0.06 | 0.96 | 1.00 | -0.01 | 0.08 | 0.08 |
| -0.52 | 1.00 | 0.01 | 0.07 | 0.07 | 0.98 | 1.00 | 0.01 | 0.04 | 0.04 |
| -0.51 | 1.00 | 0.01 | 0.04 | 0.05 | 1.01 | 1.00 | -0.01 | 0.08 | 0.08 |
| -0.50 | 1.00 | 0.01 | 0.05 | 0.05 | 1.08 | 1.00 | 0.00 | 0.05 | 0.06 |
| -0.50 | 1.00 | 0.01 | 0.04 | 0.04 | 1.10 | 1.00 | 0.00 | 0.05 | 0.05 |
| -0.49 | 0.99 | 0.00 | 0.11 | 0.11 | 1.13 | 1.00 | 0.01 | 0.04 | 0.04 |
| -0.47 | 1.00 | 0.01 | 0.05 | 0.05 | 1.14 | 1.00 | 0.01 | 0.04 | 0.04 |
| -0.45 | 1.00 | 0.01 | 0.09 | 0.09 | 1.16 | 1.00 | -0.01 | 0.07 | 0.07 |
| -0.42 | 0.99 | -0.01 | 0.13 | 0.13 | 1.22 | 1.00 | 0.01 | 0.04 | 0.04 |
| -0.33 | 1.00 | 0.01 | 0.07 | 0.07 | 1.23 | 1.00 | -0.02 | 0.09 | 0.09 |
| -0.28 | 1.00 | 0.00 | 0.09 | 0.09 | 1.43 | 1.00 | 0.00 | 0.04 | 0.04 |
| -0.27 | 1.00 | 0.01 | 0.07 | 0.07 | 1.45 | 1.00 | 0.01 | 0.04 | 0.04 |
| -0.23 | 1.00 | 0.00 | 0.09 | 0.09 | 1.47 | 1.00 | 0.00 | 0.04 | 0.04 |
| -0.21 | 1.00 | 0.01 | 0.07 | 0.07 | 1.55 | 1.00 | -0.01 | 0.07 | 0.08 |
| -0.18 | 1.00 | 0.00 | 0.10 | 0.10 | 1.60 | 1.00 | 0.01 | 0.03 | 0.03 |
| -0.15 | 0.99 | -0.01 | 0.11 | 0.11 | 1.61 | 1.00 | 0.00 | 0.05 | 0.05 |
| -0.15 | 0.99 | -0.00 | 0.11 | 0.11 | 1.62 | 1.00 | 0.00 | 0.05 | 0.05 |
| -0.11 | 1.00 | 0.01 | 0.06 | 0.06 | 1.89 | 1.00 | 0.01 | 0.03 | 0.03 |
| -0.09 | 1.00 | 0.00 | 0.09 | 0.09 | 1.91 | 1.00 | 0.01 | 0.03 | 0.03 |
| -0.05 | 1.00 | 0.01 | 0.04 | 0.04 | 1.95 | 1.00 | 0.01 | 0.02 | 0.02 |
| -0.05 | 1.00 | 0.01 | 0.04 | 0.04 | 2.25 | 1.00 | 0.00 | 0.04 | 0.04 |

We evaluated whether the model where bacteria have the associations to the PD patients is better than the model without the associations in terms of marginal likelihood. We note that the marginal likelihood represents the model evidence which expresses the preference of the data for different models. Let *M*_1_ be the model which is described Eq. 1 and *M*_0_ be the model setting all *β*_*mk*_ = 0 in Eq. 1. Table 3 shows that the marginal likelihood of *M*_1_ is greater than *M*_0_. It is better to explain the data by considering the association between the microbiota and PD.

**Table 3.** The comparison marginal likelihood.

|  | Finland | German | USA |
| --- | --- | --- | --- |
| $\mathcal{M}_0$ | -442734.62 | -5913441.14 | -3010279.35 |
| $\mathcal{M}_1$ | -355079.50 | -3807297.76 | -2063932.02 |

Figure 4 shows the estimated probabilities of the occurrences of bacteria for the three latent classes, ***p****′_l_*, (*l* = 1, 2, 3). Bacteria detected in less than three countries were removed. Arumugam *et al.* [3] showed that enterotype is characterized by the differences in the abundance of *Bacteroides, Prevotella*, and *Ruminococcus*. The result of ENIGMA shows the same tendency as previous survey. According to the results of ENIGMA, the abundance of *Enterobacteriaceae* and *Lachnospiraceae* also differ greatly among clusters. Bacterial abundance differs between countries. In USA there is a large abundance of *Verrucomicrobiaceae*, but in Finland there are few. Conversely, in Finland there is more *Prevotellaceae*, but in USA it is less.

**Figure 4.**
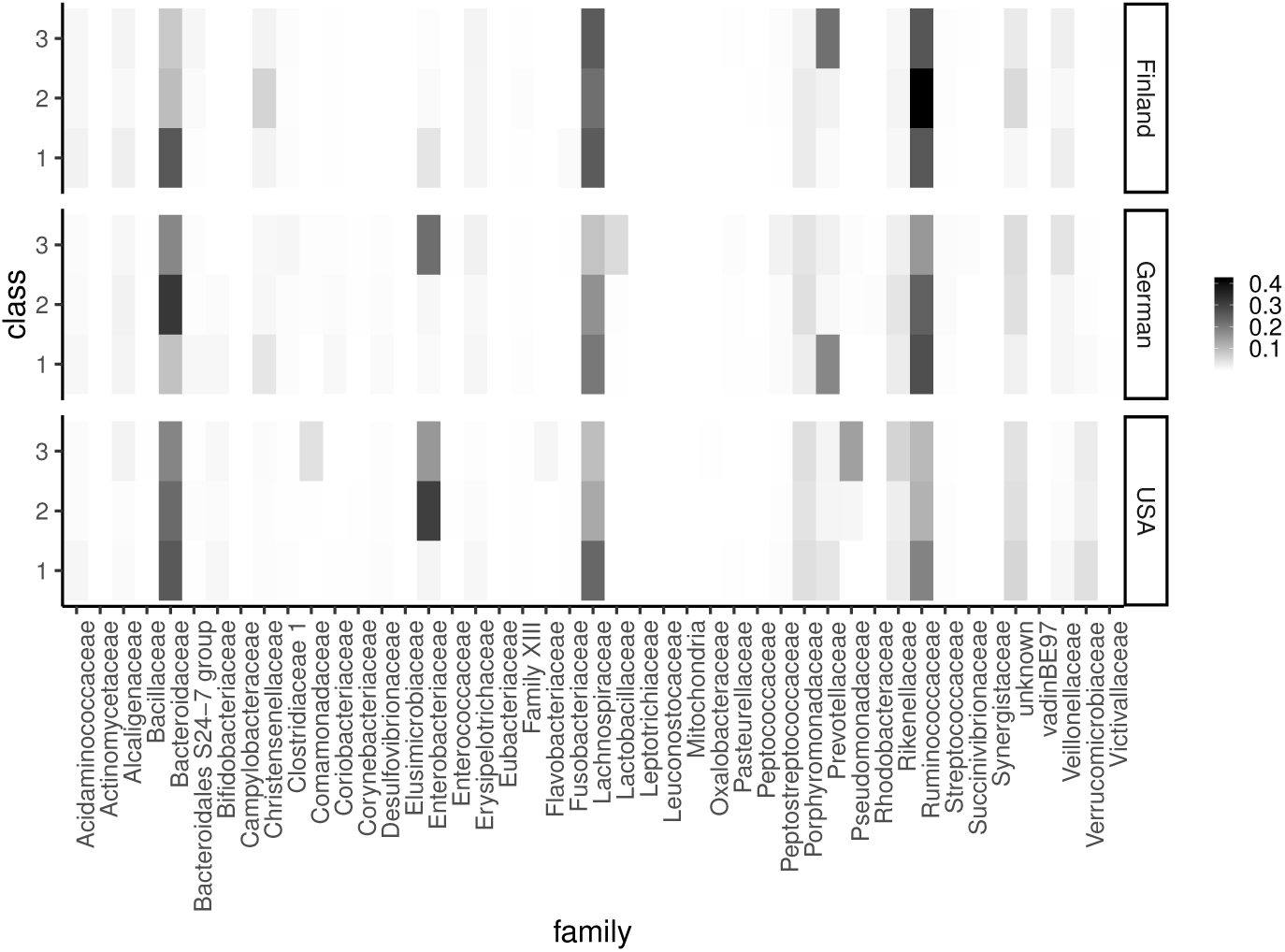
Heatmap showing 1 (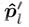). This quantities corresponds to the probabilities of the occurrences of bacteria for the three latent classes.

Table 5 shows the coefficients whose 95% credible intervals do not contain zero in more than two countries. The microbes with these coefficients indicates that the corresponding microbial composition patterns are significantly related to PD. We found that, in family levels, *Clostridiaceae, Comamonadaceae, Pasteurellacea, Prevotellaceae, Actinomycetaceae, Bifidobacteriaceae, Enterococcaceae, Lactobacillaceae, Synergistaceae, Verrucomicrobiaceae* and *Victivallaceae*, the signs of these coefficients matched in all countries. These results are consistent with previous studies. Hill-Burns *et al.* [17] reported PD patients contained high levels of *Bifidobacteriaceae* and *Verrucomicrobiaceae* and low levels of *Pasteurellaceae*. *Scheperjans et al.* [16] reported PD patients contained high levels of *Lactobacillaceae, Verrucomicrobiaceae* and low levels of *Prevotellaceae*. Hopfner *et al* reported PD patients have high levels of *Lactobacillaceae* and *Enterococcaceae*.

We compared ENIGMA to the Wilcoxon rank sum test, one of the classical methods for identifying bacteria related with a environmental factor of interest [18]. Table 4 shows bacteria significantly related with the PD patients with *p*-value < 0.05 in more than two countries. We observed that the bacteria detected by the Wilcoxon test were almost included in those of ENIGMA (Table 5). We note that all of the corrected *p*-values in Table 4 are larger than 0.05. This result shows that ENIGMA is superior to the Wilcoxon rank sum test in terms of identifying more associations between microbiota and the PD patients.

**Table 4.**
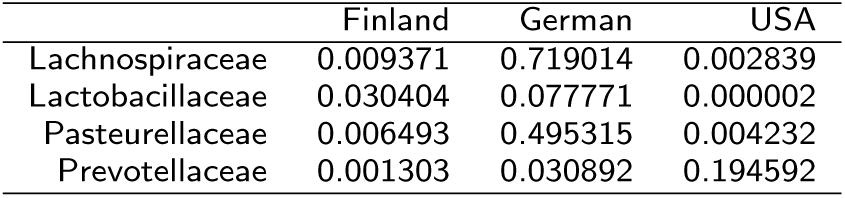
p-value of Wilcoxon test Abe *et al.*

|  | Finland | German | USA |
| --- | --- | --- | --- |
| Lachnospiraceae | 0.009371 | 0.719014 | 0.002839 |
| Lactobacillaceae | 0.030404 | 0.077771 | 0.000002 |
| Pasteurellaceae | 0.006493 | 0.495315 | 0.004232 |
| Prevotellaceae | 0.001303 | 0.030892 | 0.194592 |

**Table 5.** The bacteria which significant associated with PD in more than two countries. The “-” notation indicates the bacteria undetected in that country.

| family | Finland |  |  | German |  |  | USA |  |  |
| --- | --- | --- | --- | --- | --- | --- | --- | --- | --- |
| | $\hat{\beta}$ | lower bound | upper bound | $\hat{\beta}$ | lower bound | upper bound | $\hat{\beta}$ | lower bound | upper bound |
| Anaeroplasmataceae | -0.87 | -1.28 | -0.45 | -1.69 | -2.03 | -1.35 | - | - | - |
| Bacteroidales S24-7 group | -0.52 | -0.93 | -0.11 | 0.22 | -0.12 | 0.56 | -0.69 | -1.04 | -0.33 |
| Bradyrhizobiaceae | - | - | - | -0.82 | -1.17 | -0.47 | -1.51 | -2.29 | -0.74 |
| Brevibacteriaceae | - | - | - | -1.02 | -1.38 | -0.66 | -0.58 | -0.97 | -0.19 |
| Brucellaceae | - | - | - | -1.69 | -2.50 | -0.87 | -1.35 | -1.76 | -0.94 |
| Clostridiaceae 1 | -0.54 | -0.96 | -0.13 | -0.08 | -0.42 | 0.26 | -0.43 | -0.79 | -0.08 |
| Comamonadaceae | -0.85 | -1.35 | -0.35 | -1.27 | -1.61 | -0.93 | -0.20 | -0.55 | 0.16 |
| Elusimicrobiaceae | -4.17 | -5.60 | -2.74 | -2.11 | -2.54 | -1.68 | 2.36 | 1.04 | 3.67 |
| Intrasporangiaceae | - | - | - | -3.47 | -4.86 | -2.07 | -3.03 | -4.77 | -1.28 |
| Leuconostocaceae | -2.66 | -4.30 | -1.02 | 0.50 | 0.13 | 0.86 | -1.63 | -2.11 | -1.15 |
| Moraxellaceae | - | - | - | -1.58 | -1.92 | -1.24 | -0.91 | -1.26 | -0.56 |
| Pasteurellaceae | -1.62 | -2.07 | -1.17 | 0.30 | -0.04 | 0.64 | -1.80 | -2.16 | -1.44 |
| Prevotellaceae | -2.46 | -2.87 | -2.05 | -0.03 | -0.37 | 0.30 | -0.45 | -0.80 | -0.09 |
| Rhodocyclaceae | - | - | - | -3.53 | -4.93 | -2.13 | -0.70 | -1.13 | -0.27 |
| Actinomycetaceae | 0.11 | -0.78 | 1.01 | 0.42 | 0.07 | 0.78 | 1.07 | 0.70 | 1.43 |
| Bacillaceae | 1.72 | 0.34 | 3.11 | -2.35 | -2.72 | -1.99 | 0.86 | 0.50 | 1.22 |
| Bdellovibrionaceae | - | - | - | 1.43 | 0.40 | 2.46 | 2.87 | 1.69 | 4.05 |
| Bifidobacteriaceae | 1.34 | 0.82 | 1.86 | 0.54 | 0.20 | 0.88 | 0.09 | -0.26 | 0.45 |
| Campylobacteraceae | 0.36 | -0.31 | 1.03 | 4.90 | 4.48 | 5.33 | 1.04 | 0.67 | 1.41 |
| Cytophagaceae | - | - | - | 2.45 | 1.56 | 3.34 | 1.50 | 0.20 | 2.81 |
| Enterococcaceae | 3.87 | 2.70 | 5.05 | 0.74 | 0.40 | 1.08 | 0.16 | -0.20 | 0.52 |
| Lactobacillaceae | 3.00 | 2.56 | 3.43 | -0.51 | -0.85 | -0.18 | 1.80 | 1.44 | 2.16 |
| Leptotrichiaceae | -0.90 | -1.89 | 0.09 | 2.57 | 1.88 | 3.26 | 0.92 | 0.46 | 1.37 |
| Methanobacteriaceae | - | - | - | 0.93 | 0.59 | 1.27 | 0.76 | 0.39 | 1.12 |
| Mitochondria | 0.60 | -1.27 | 2.46 | 0.73 | 0.11 | 1.36 | 1.60 | 0.98 | 2.21 |
| Paenibacillaceae | - | - | - | 2.19 | 1.28 | 3.10 | 1.73 | 1.32 | 2.13 |
| Planococcaceae | - | - | - | 1.06 | 0.72 | 1.41 | 3.26 | 2.69 | 3.84 |
| Rhizobiaceae | - | - | - | 0.64 | 0.24 | 1.03 | 1.52 | 1.09 | 1.94 |
| Streptococcaceae | 0.44 | 0.03 | 0.86 | 0.84 | 0.50 | 1.17 | 0.34 | -0.02 | 0.69 |
| Succinivibrionaceae | -0.32 | -0.76 | 0.11 | 0.74 | 0.40 | 1.08 | 4.29 | 3.76 | 4.82 |
| Synergistaceae | 1.26 | 0.80 | 1.71 | 0.25 | -0.10 | 0.61 | 1.51 | 1.14 | 1.89 |
| Verrucomicrobiaceae | 1.71 | 1.23 | 2.19 | 1.62 | 1.29 | 1.96 | 0.03 | -0.32 | 0.39 |
| Victivallaceae | 0.42 | -0.00 | 0.85 | 0.68 | 0.34 | 1.02 | 1.02 | 0.63 | 1.40 |

The analyses with real data thus show that ENIGMA can identify enterotype-like clusters and the associations between the gut microbiota and the PD patients, and some of the results are strongly supported by the previous researches.

## Conclusion

We proposed a novel hierarchical Bayesian model, ENIGMA, to discover the underlying microbial community structures and associations between microbiota and their environmental factors from microbial metagenome data. ENIGMA is based on a probabilistic model of a microbial community structures and supplied with labels for one or more environmental factors of interest for each sample. The structures of each sample is modeled by a multinomial distribution whose parameters are represented independently by group and environmental effects of each sample, which prevent mixing of individual differences and effects of interest. This framework enables the model to learn (*i*) how microbes contribute to an underlying community structures (cluster) and (*ii*) how microbial compositional patterns are explained environmental factors of interest, simultaneously. The effectiveness of ENIGMA was evaluated on the bases of experiments involving both synthetic and read datasets. We believe that these newly discovered clusters and associations estimated from ENIGMA would provide more insight in the the mechanisms of a microbial community.

There is one major limitation of ENIGMA is its scalability and efficiency, since the number of the parameters in the model grow proportional to the number of taxa when the number of environmental factors of interest is large. Further works should focus on developing a scalable probabilistic model of microbial compositions to analyze underlying microbial structures with a large number of these effects by using sparse parameter estimation [20]. We are also interested in developing a dynamic probabilistic model similar to reproted by Blei and Lafferty [21] to analyze time-varying bacteria compositions during the progression of a disease.

## Author’s contributions

KA and TS designed the proposed algorithm; KO and MH designed the experiments.

## Acknowledgements

This work was supported by Grants-in-Aid from the Ministry of Education, Culture, Sports, Science and Technology of Japan (MEXT); Ministry of Health, Labour and Welfare of Japan (MHLW); Japan Agency for Medical Research and Development (AMED), and the Hori Sciences and Arts Foundation.

